# Rice Bran and Quercetin Produce a Positive Synergistic Effect on Human Gut Microbiota, Elevate the Level of Propionate, and Reduce the Population of *Enterobacteriaceae* family when Determined using a Bioreactor Model

**DOI:** 10.1101/2020.02.13.947515

**Authors:** Sudeep Ghimire, Supapit Wongkuna, Ranjini Sankaranarayanan, Elizabeth P. Ryan, G. Jayarama Bhat, Joy Scaria

## Abstract

Diet is one of the prominent determinants of gut microbiota composition significantly impacting human health. Recent studies with dietary supplements such as rice bran and quercetin have been shown to provide a beneficial impact on the host by positively influencing the gut microbiota. However, the specific bacterial species impacted when rice bran or quercetin is present in the diet is not well understood. Therefore, in this study, we used a minibioreactor array system as a model to determine the effect of quercetin and rice bran individually, as well as in combination, on gut microbiota without the confounding host factors. We found that rice bran exerts higher shift in gut microbiome composition when compared to quercetin. At the species level, *Acidaminococcus intestini* was the only significantly enriched taxa when quercetin was supplemented, while 15 species were enriched in rice bran supplementation and 13 were enriched when quercetin and rice bran were supplemented in combination. When comparing the short chain fatty acid production, quercetin supplementation significantly enriched isobutyrate production while propionate dominated the quercetin and rice bran combined group. Higher levels of propionate were highly correlated to the lower abundance of the potentially pathogenic *Enterobacteriaceae* family. These findings suggest that the combination of rice bran and quercetin serve to enrich beneficial bacteria and reduce potential opportunistic pathogens. However, further *in vivo* studies are necessary to determine the synergistic effect of rice bran and quercetin on host health and immunity.

**Importance:** Rice bran and quercetin are dietary components that shape host health by interacting with the gut microbiome. Both these substrates have been reported to provide nutritional and immunological benefits individually. However, considering the complexity of the human diet, it is useful to determine how the combination of food ingredients such as rice bran and quercetin influences the human gut microbiota. Our study provides insights into how these ingredients influence microbiome composition alone and in combination *in vitro*. This will allow us to identify which species in the gut microbiome are responsible for biotransformation of these dietary ingredients.. Such information is helpful for the development of synbiotics to improve gut health and immunity.

## Introduction

Gut microbiome influences health and disease (1-3) and is highly affected by diet (4-6). Some of the dietary components are known to enhance host health by promoting the growth of beneficial bacteria in the gut, termed as “prebiotics” (7, 8). Rice bran and quercetin have been used as prebiotics in mice and pig experiments, showing a significant improvement of the gut microbiota (9-11). Rice bran is a byproduct of the rice milling process and is an easily available, affordable, sustainable, and globally produced source of prebiotics. It is known to supplement nutrients (12), increase beneficial bacteria growth and enhance gut mucosal immunity (10), and prevent diseases (9, 13-15). Similarly, quercetin (3,3′,4′,5,7-pentahydroxyflavone) represents an important subgroup of flavonoids found in fruits and leafy vegetables (16). It has received substantial attention in the past few years from the scientific community due to its anti-inflammatory effects (17, 18) and its potential health-promoting properties in the treatment or prevention of cardiovascular diseases (19), lung and colorectal cancers (20, 21), and colitis (11, 22). Furthermore, the mammalian body can endure high levels of quercetin without any significant adverse health effects (23) suggesting its potential food application.

Despite multiple studies corroborating the beneficial effect of rice bran and quercetin on the host, their effect on microbiota varies significantly from study to study and has lower taxonomic resolution (22, 24-26). Variation in *in vivo* studies are due to multiple compounding host-related factors which makes it difficult to interpret microbiome results precisely (27). In contrast, studying the effect of dietary ingredients on the microbiome in the absence of confounding host factors can help to better understand the microbial interactions. Minibioreactor array is an *in vitro* model system that supports the growth of complex, stable microbiota which mimic hindgut conditions without the complexity of host factors(28). Bioreactors have been used as model systems to study how dietary ingredients shape microbiome(29). The use of bioreactors for such studies allows precise control of the environmental conditions that affect microbiome composition which provide increased reproducibility and reveal microbial interactions in a more defined way. Furthermore, it would also help to identify microbial biotransformations and metabolites more precisely(30). We used minibioreactor array systems to gain deeper understanding of how the microbiota responds to rice bran or quercetin without host interference. Since supplementation of rice bran and quercetin individually have shown positive effects on host health, we hypothesized that a combination of rice bran and quercetin will have a synergistic positive effect on the gut microbiota. Using minibioreactors as a model (28, 31), we sought to understand the impact of rice bran and quercetin individually and in combination on the human gut microbiome. The bioreactors were operated continuously for 21 days and samples were collected for 16S rRNA sequencing, metagenomic sequencing and short chain fatty acid analysis. We observed that the rice bran and quercetin combination was effective in significantly reducing *Enterobacteriaceae* family members and had higher propionate levels. This study provides insights of species-level shifts in gut microbiome in the presence of quercetin and/or rice bran supplementation through an *in vitro* model.

## Materials and Methods

### Donor samples, mini-bioreactor array preparation and sample collection

We obtained fresh fecal samples from six healthy donors with no prior history of antibiotic consumption in the past year. The six fecal samples were pooled in equal proportion (32) and used as inoculum. The modified BHI medium (32) was used as a control medium. Rice bran, originating from rice variety RBT 300 (USA), was prepared as described previously (12) and added to the control medium (final concentration: 2 mg/mL): designated as RB. Quercetin was added to the media at a final concentration of 75 mg/mL and is referred to as QC. The final experimental group consisted of modified BHI medium with the additions of quercetin (75 mg/mL) and rice bran extract (2 mg/mL) and is identified as QC+RB. Mini-bioreactors (MBRAs) were sterilized, assembled and the experiment was performed as described previously (31) with minor modifications. Briefly, the input and output on Watson Marlow pumps were set at 1 rpm and 2 rpm respectively. The rotating magnetic stirrer was set at 130 rpm. The media (Control, QC, RB, and QC+RB) (Figure 1A) were allowed to flow continuously for 24 hours each in triplicate. Three hundred microliters of the inoculum was introduced into all wells with a retention time of 16 hours. The continuous flow model was operated up to 21 days post-inoculation (Figure 1B). Five hundred microliters of the media was collected for sequencing at day 0 (inoculum), days 4, 7, 14 and 21 and directly frozen to −80°C. Also, samples for short chain fatty acid (SCFAs) determination were collected at days 4, 7, 14 and 21 post-inoculation (Figure 1B).

**Figure 1:**
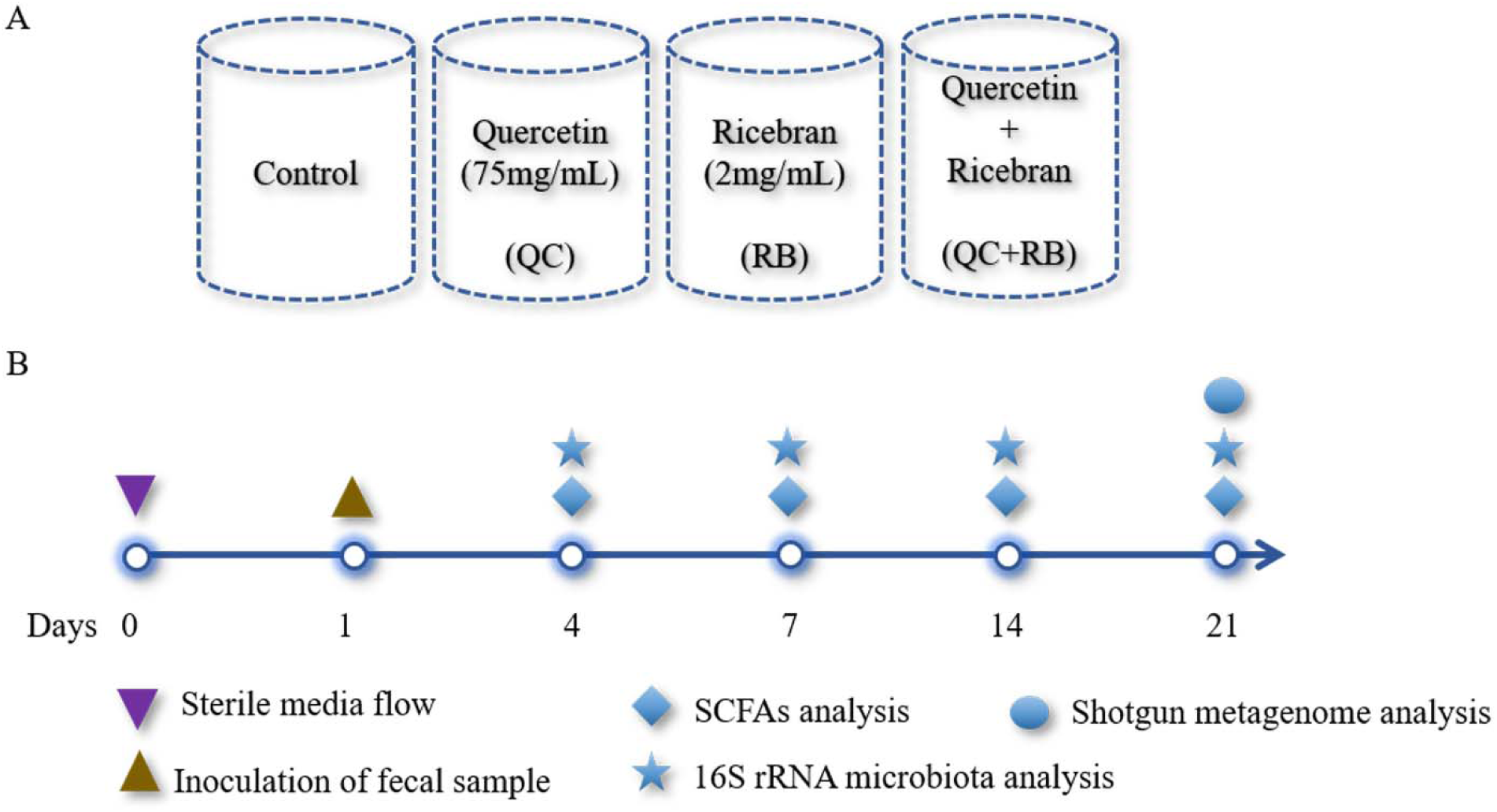
Overview of the study design: **A)** A schematic diagram shown the design of four different media conditions used in this study. Single substrates (quercetin or rice bran) or the mixture of quercetin and rice bran were used at the final concentration of 75 mg/L and 2 mg/mL as shown above in the base medium BHI (see Methods section). Final pH of the media was adjusted to 6.8±0.2 for all conditions. Each condition was run in triplicate and were inoculated with the same fecal inoculum (see Methods section). **B)** Outline of the bioreactor experiment showing time points for fecal inoculation and sample collection for SCFAs and microbial community analysis.

### Microbial DNA extraction and sequencing

DNA isolation was performed on 50 samples including duplicate inoculum samples. The DNA was extracted from 500 µl of the sample using a Powersoil DNA isolation kit (MoBio Laboratories Inc, CA) following the manufacturer’s instructions. After extraction, the quality of DNA was measured using NanoDrop™ one (Thermo Fisher Scientific, DE) and quantified using Qubit Fluorometer 3.0 (Invitrogen, CA). The DNA samples were stored at −20°C until further use. To analyze the variation of the microbial composition over time, all samples were amplicon sequenced using an Illumina MiSeq platform with paired-end V3 chemistry. The library was prepared using an Illumina Nextera XT library preparation kit (Illumina Inc, CA) targeting V3-V4 regions of the 16S rRNA. The libraries were bead normalized and multiplexed before loading into the sequencer. We also performed shotgun metagenome sequencing on twelve samples obtained from day 21 to identify species level taxonomical differences between the experimental groups. We used Nextera XT kit (Illumina, San Diego, CA) for the preparation of the shotgun metagenome sequencing library. The library was then sequenced using paired end 300 base sequencing chemistry using MiSeq platform.

### Data analysis

The time-series changes in the microbial communities were analyzed using 16S rRNA community analysis in Quantitative Insights into Microbial Ecology framework (QIIME, Version 2.0) (33). Briefly, the demultiplexed reads obtained were quality filtered using q2-demux plugin and denoised applying DADA2 (34). All amplicon sequence variants were aligned with *mafft* (35) to construct a phylogeny with fasttree2 (36). The outputs rooted-tree.qza, table.qza, taxonomy.qza were then imported into R (37) for analysis using *phyloseq* (38). Shannon diversity and bray curtis dissimilarity indices were calculated as alpha and beta diversity metrices. Kruskal-Wallis test at p=0.05 was performed to compare the species richness between the groups. The reads were normalized by rarefying to 30,000 and the taxonomy was assigned to amplicon sequence variants using the q2-feature-classifier (39) using Greengenes as the reference (40). Rarefying the reads to 30,000 were enough to estimate total diversity and taxonomy (Supplementary Figure 1A). Initially, a total of 947 amplicon variants were identified from 50 samples. The average number of non-chimeric reads per sample obtained in Qiime2 pipeline was 92,512±27,631 (mean ± SD). The amplicon sequence variants obtained were filtered to select those which are present in at least 20% of samples with a count of 10 each or amplicon sequence variants with > 0.001% of total median count reducing the total number to 581.

For shotgun metagenomes, raw fastq sequences were quality controlled using Fastqc (https://www.bioinformatics.babraham.ac.uk/projects/fastqc/) and host reads were removed using metaWRAP pipeline (41). Filtered reads were analyzed for taxonomy using Kaiju (42) against the proGenomes database (43) using default parameters. The percentage abundance of each taxon was plotted using Explicit v2.10.5 (44). The samples from day 21 yielded a total of 39,332,649 reads of which 70. 86% were classified to 161 different species, whereas 185,188 (5.55%) and 9,278,647 (23.59%) reads remained as unclassified bacteria or unassigned to non-viral species, respectively. In addition, the raw counts obtained for each taxon were used for calculation of alpha and beta diversity using the *phyloseq* package in R. For identification of differentially abundant taxa in QC, RB and QC+RB compared to control medium, the raw counts of abundance of each taxon was then Log10(x+1) transformed and fed to DESEq2 (45) package in R. Bacterial taxa which significantly altered from the control were further filtered out with the criteria of at least log2foldchange (Log2FC) of ≥ 2 and p*adj* value > 0.05. The enriched taxa were selectively analyzed for their co-relation to SCFAs phenotypic data using Spearman correlation method in R.

### Estimation of Short Chain Fatty Acids (SCFAs)

For the estimation of SCFAs, 800 µl of samples were collected from each mini-bioreactor, mixed with 160 µl of 25% m-phosphoric acid and frozen at −80°C until further analysis. Later, the frozen samples were thawed and centrifuged (>15,000 × g) for 20 min. Five hundred microliters of supernatant was collected in the tubes before loading into the gas chromatograph (Agilent Technologies, USA) for analysis. The SCFA concentrations were compared between the groups using the Kruskal-Wallis test followed by the Dunn test with Benjamini-Hochberg correction in R and visualized using GraphPad Prism 6.0

## RESULTS

### Rice bran caused higher shift in bacterial population when compared to quercetin

The microbial community richness indicated by the Shannon diversity index showed significant differences in the early days (days 4 and 7) due to addition of rice bran in the media whether alone or in combination with quercetin (Supplementary Figure 1B). No significant changes in the richness was observed among the groups on day 14 and day 21, suggesting the stabilization of the microbial communities by day 14. Also, on day 21, shotgun metagenome analysis showed no differences in the richness between the four conditions (Figure 2B). In contrast, marked differences between the communities were evident as early as day 4 by beta diversity analysis (Supplementary Figure 2A). Rice bran supplementation showed greater shift in the bacterial community as shown on days 7, 14 and 21 (Supplementary Figure 2B, 2C, 2D). Based on Bray-Curtis distance, RB and QC+RB treatments were similar to one another and clustered separately from control and QC. This suggest that the addition of quercetin made limited shift in the community compared to rice bran (Supplementary Figure 2). Shotgun metagenome sequencing at day 21 also indicated significant differences in the community profile, primarily influenced by the addition of rice bran extract in the medium (Figure 2C).

**Figure 2:**
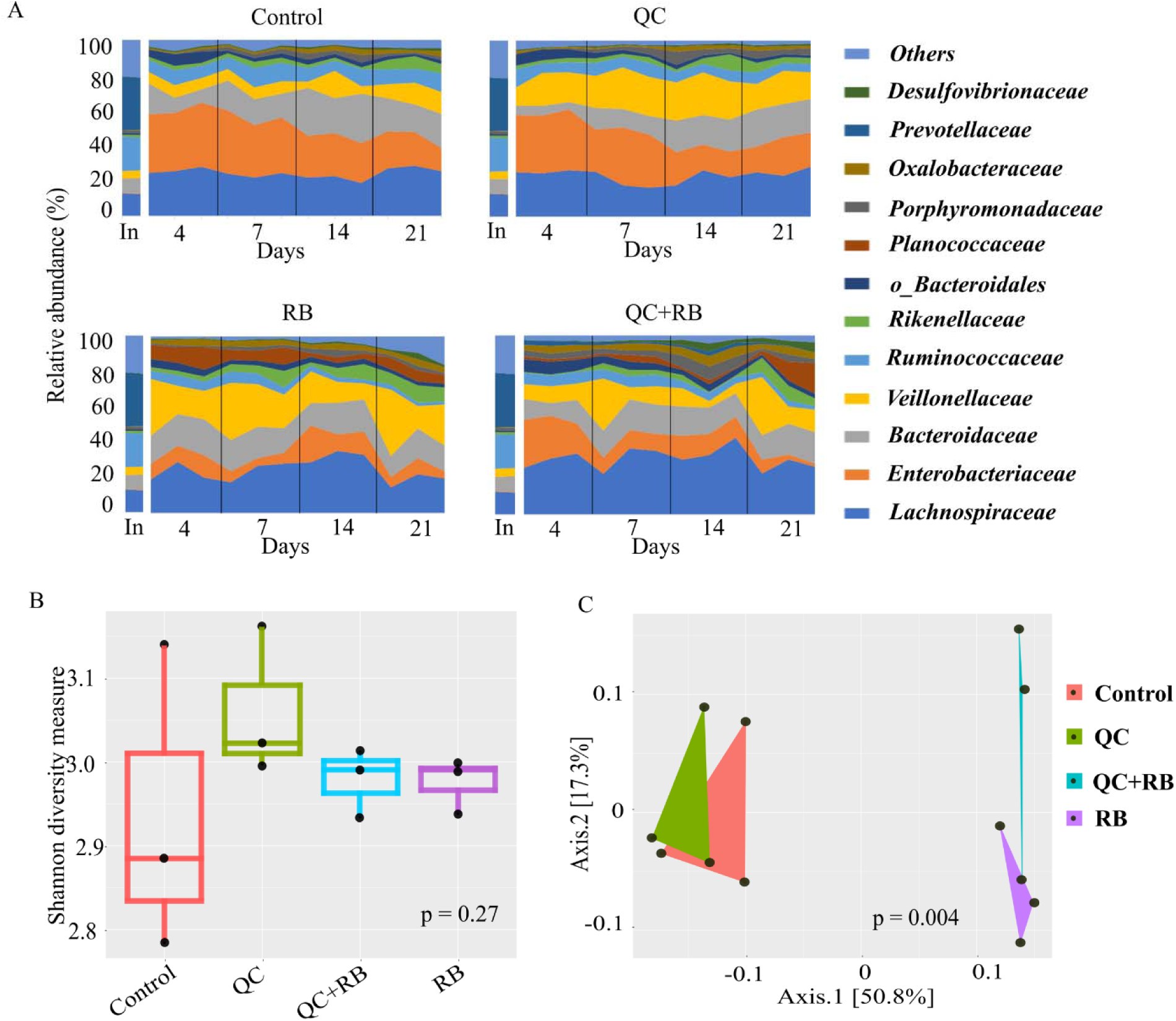
Gut microbiota compositional changes following rice bran and quercetin supplementation. **A)** Temporal family-level composition of the most abundant 12 bacterial taxa at day 4, 7, 14 and 21 and inoculumdetermined using 16S rRNA community profiling for Control, QC, RB and QC+RB conditions. **B)** Alpha diversity measure (Shannon diversity) between Control, supplemented with QC, supplemented with RB and supplemented with QC+RB determined using shotgun metagenome sequence analysis at day 21. Kruskal-Wallis test was performed for the Shannon diversity indices obtained for the groups (p=0.27). **C)** Beta diversity among the control, QC, RB and QC+RB groups were obtained using shotgun metagenome sequence analysis on day 21. “MDS” ordination followed by Bray-Curtis distance calculation was used to visualize the differences between the groups (adonis, p= 0.004).

When examined taxonomically, the inoculum was dominated by *Prevotellaceae* (29.9±0.87 %), followed by *Ruminococcaceae* and *Lachnospiraceae*. However, the abundance of *Prevotellaceae* was reduced to ∼ 0.0 % in all groups suggesting that not all taxa in the inoculum were supported for growth. Over time, there was a major shift in the composition of the microbiota between the groups starting as early as day 4 (Figure 2A). *Enterobacteriaceae* dominated the control and quercetin groups on day 4, followed by *Lachnospiraceae*. However, by day 21, *Lachnospiraceae* dominated the control and QC conditions followed by *Bacteroidaceae* and *Enterobacteriaceae*. In medium supplemented with rice bran extract (RB), *Lachnospiraceae* was dominant followed by *Veillonellaceae* and *Bacteroidaceae* on day 4. By day 21, *Veillonellaceae* was the dominant family. *Lachnospiraceae* also dominated the community in the quercetin and rice bran combination (QC+RB) early on, followed by *Enterobacteriaceae*. On day 21, *Lachnospiraceae* still dominated followed by *Veillonellaceae* and *Bacteroidaceae*. Even though *Lachnospiraceae* was dominant in the control, QC and QC+RB medium at day 21, *Enterobacteriaceae* was highly reduced only for QC+RB.

On day 21, we performed shotgun metagenome analysis to identify the species level differences among the four groups (Figure 3). Based on DeSEQ2 analysis, 13 species were altered significantly in the quercetin group (Supplementary Figure 3A), 70 species in the rice bran group (Supplementary Figure 3B) and 53 species in the quercetin and rice bran combination group compared to the control (Figure 4). With a cut off Log2FC≥ 2 and p*adj* value > 0.05, *Acidaminococcus intestine* was the only enriched taxa in media supplemented with quercetin (QC) while *Coprococcus catus, Dorea longicatena, Anaerostipes caccae, Eubacterium limosum* and *Enterococcus faecalis* were highly reduced. Interestingly, the population of *Flavonifractor plautii*, a flavonoid metabolizing bacterium (46, 47) was significantly reduced in quercetin supplementation. The populations of *F. plautii* and *Pseudoflavonifractor capillosus* also were reduced in the group supplemented with quercetin and rice bran. The combination (QC+RB) enriched 13 additional species and reduced 13 others. Rice bran supplementation alone (RB) enriched15 species including six different species from *Veillonella* (Supplementary Table 1).

**Figure 3:**
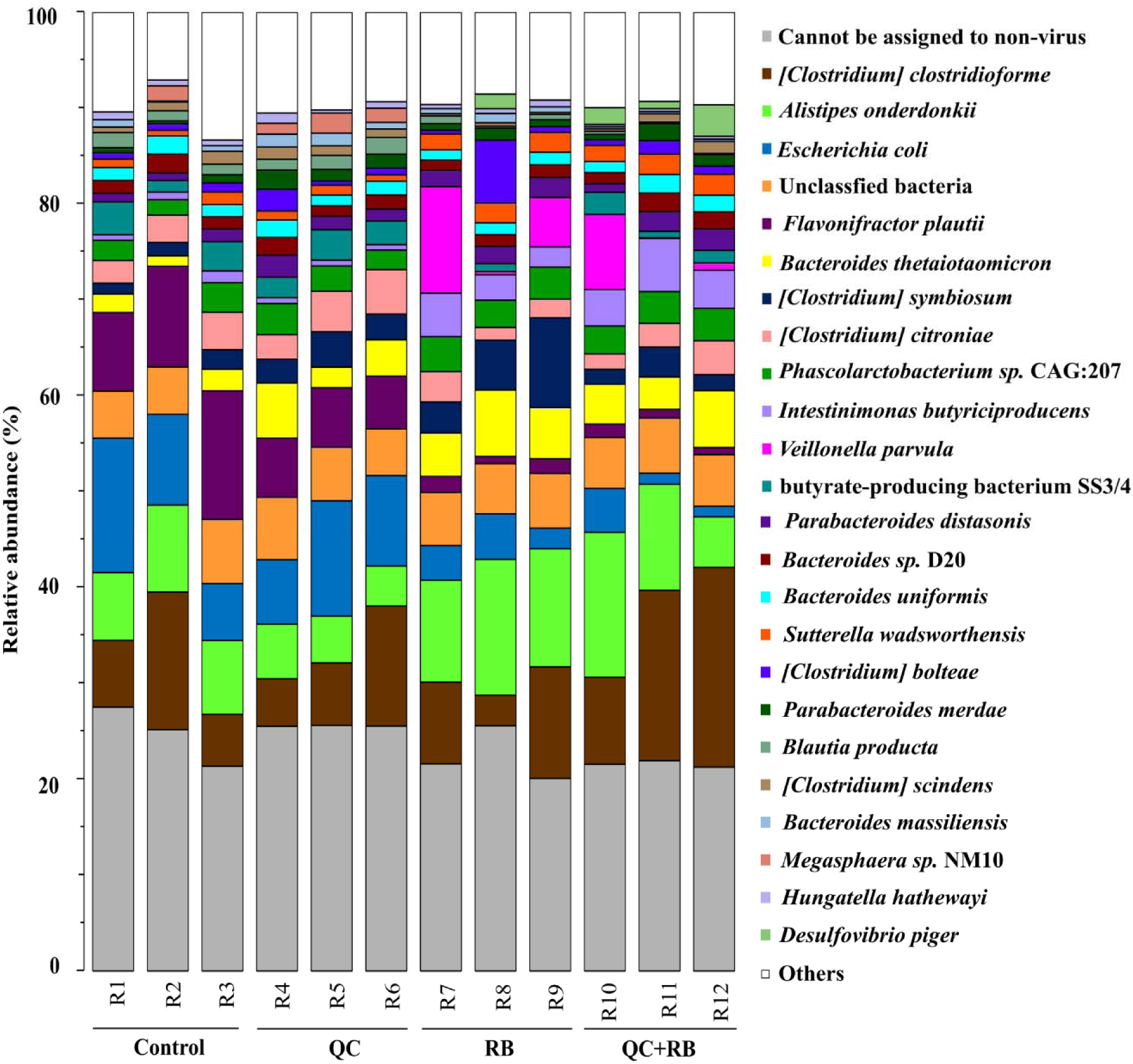
Impact of quercetin (QC) and rice bran (RB) supplementation on bacterial taxonomy. Species-wise variation of the most abundant (top 25) of the bacterial taxa determined by shotgun metagenome sequencing at day 21 (endpoint) for control, QC, RB and QC+RB conditions in triplicate.

**Figure 4:**
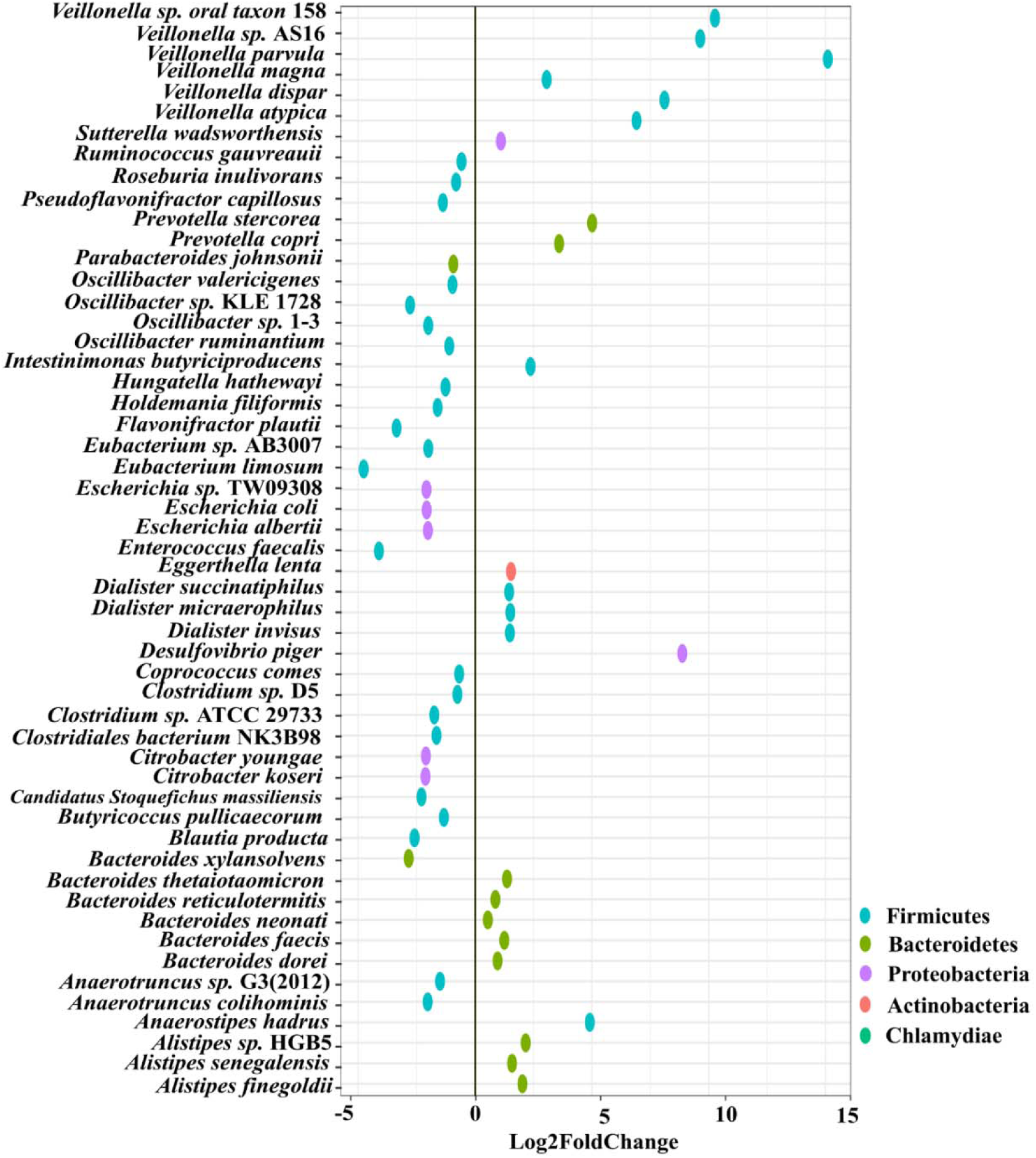
Impact of quercetin and rice bran (QC+RB) supplementation on bacterial species that are significantly altered in QC+RB when compared to the control medium. The differential significance of the species was calculated by using DESEq2 (See Supplementary Table 1) from shotgun metagenome sequence analysis at day 21. The left column represents bacterial species-level taxa while the colors indicate their corresponding phyla (“Blue” =*Firmicutes*, “Brown” =*Bacteroidetes*, “Pink” =*Proteobacteria*, “Orange” =*Actinobacteria*, “Green” =*Chlamydiae*).

### Rice bran and quercetin combination yield higher propionate levels and reduces members of *Enterobacteriaceae* family

Along with the changes in the taxonomy of the microbial community, substrate inclusion into a medium alters the metabolic profile of a community (48, 49). To understand how these substrates have altered the fermentation potential of the microbiota, we measured SCFAs from each minibioreactor on days 4, 7, 14 and 21. Acetate, propionate and butyrate were the major SCFAs produced, while isobutyrate, valerate and isovalerate were present in lower concentrations. The amount of each SCFA produced was stable by day 14 and at the endpoint, with marked differences between the groups (Figure 5A, 5B, 5C, 5D, 5E). At day 21, communities formed in the control and RB weighted strongly towards acetate and butyrate production respectively (Supplementary Figure 4). Statistically, acetate production was similar in the control and QC+RB group and significantly different from the RB group (Figure 5F). The butyrate production in the RB medium was higher but not significantly different compared to the other groups (Figures 5H). Similarly, the medium supplemented with quercetin and rice bran (QC+RB) weighted towards the production of propionate and butyrate (Supplementary Figure 4). Rice bran supplementation significantly raised propionate levels in combination with quercetin compared to the QC group (Figure 5G). In contrast, quercetin supplementation in the medium, weighted towards the production of minor SCFAs, i.e. valerate, isovalerate and isobutyrate. However, only isobutyrate production was significantly higher in QC compared to QC+RB group (Figure 5I).

**Figure 5:**
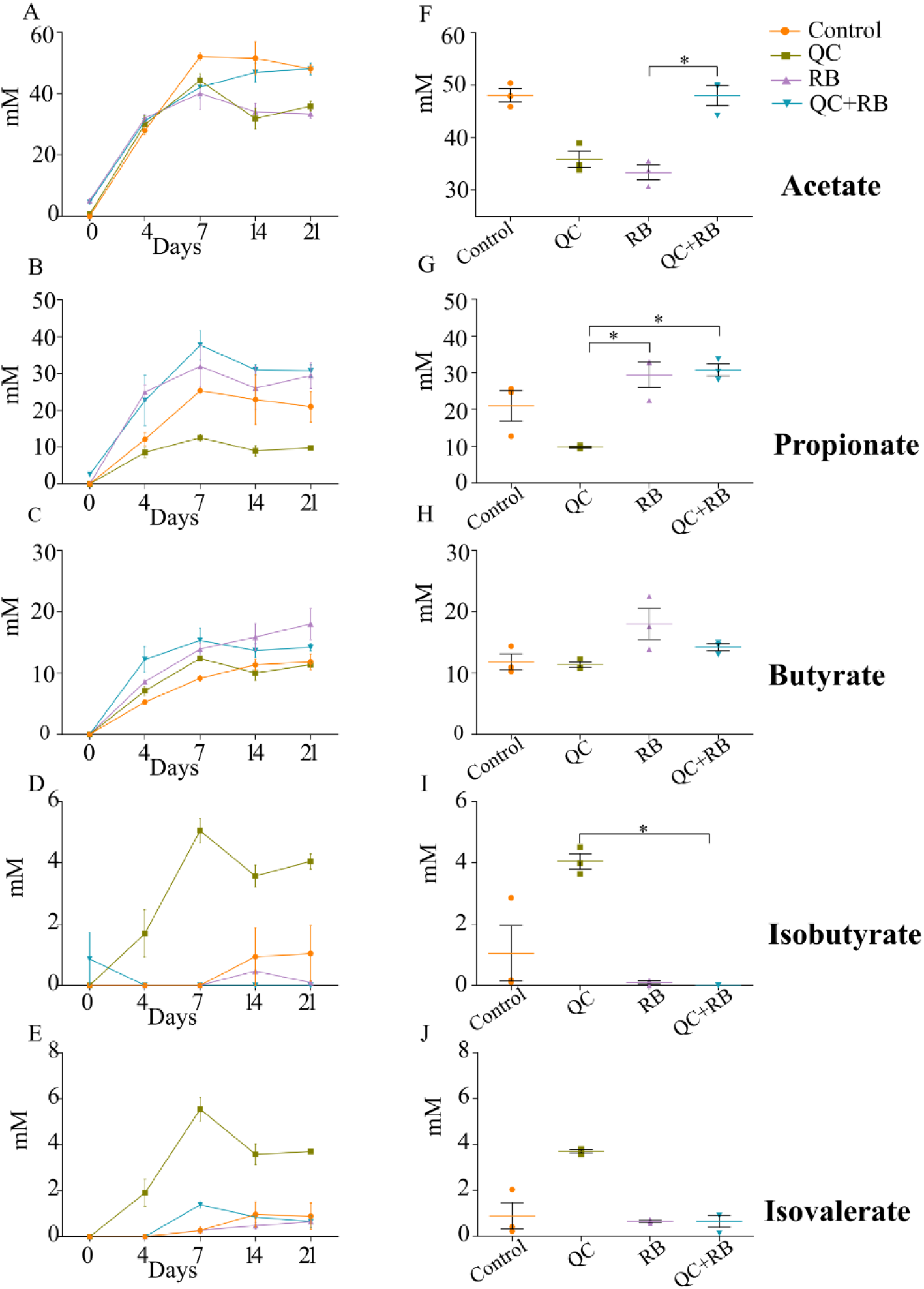
Effect of quercetin (QC) and rice bran (RB) on short Chain Fatty Acids (SCFAs) production in QC, RB, and QC+RB compared to control medium in minibioreactors. A-E represent the periodic variation of acetate, propionate, butyrate, isobutyrate and isovalerate at day 0, 4, 7, 14 and 21 respectively. F-J represent the comparison of concentrations of acetate, propionate, butyrate, isobutyrate and isovalerate respectively from control, QC, RB and QC+RB medium at day 21 (endpoint). Kruskal-Wallis test was performed between the groups and *posthoc* (Dunn test) analysis was performed to identify the different significant groups. “*” represents significance at 0.05. Error bars represent standard error of the mean of data obtained from three different bioreactors.

SCFAs production by microbiota has been associated with pathogen inhibition to benefit the host (50). Physiological levels of SCFAs are reported to reduce *Enterobacteriaceae* members by pH mediated action (51). Specifically, propionate production by a propionate producing consortium has been shown to reduce antibiotic induced dysbiosis (52). In this study, as higher production of propionate weighted towards QC+RB medium at day 21, we estimated the correlation of *Enterobacteriaceae* abundance to propionate levels. Significant high negative correlations were observed between the abundance of the members of *Enterobacteriaceae* family and propionate with media change (Figure 6). *Citrobacter rodentium, C. koseri, C. youngae, Escherichia albertii, E. coli* and *Escherichia* sp. TW09308 (Figure 6A, B, C, D, E, F) were greatly reduced in QC+RB medium, whereas propionate levels were high.

**Figure 6:**
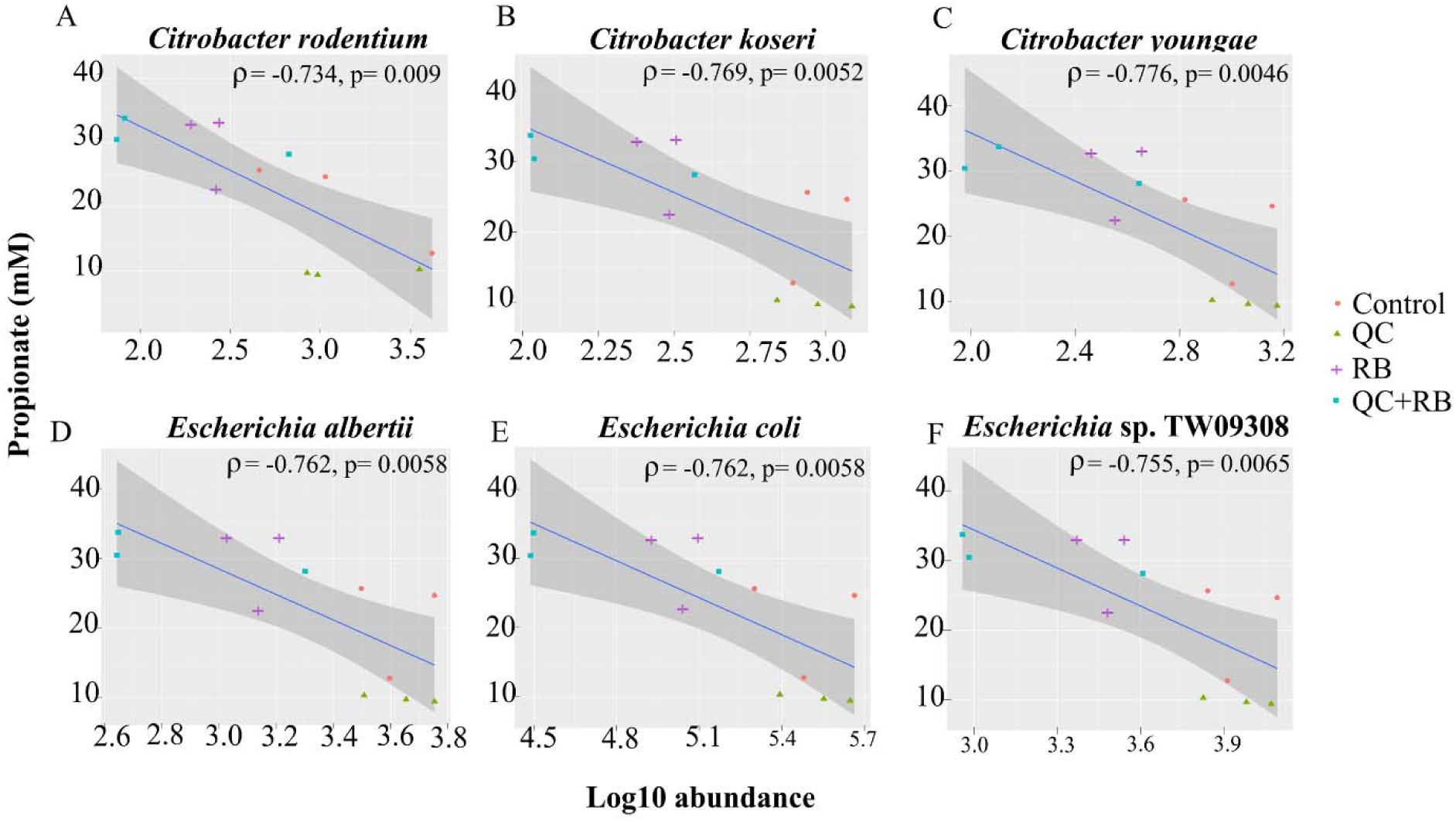
Spearman correlation of *Enterobacteriaceae* family members (*Citrobacter* and *Escherichia* species) log10 abundances at day 21 determined by shotgun metagenomics sequence analysis to levels of propionate in the medium. A negative correlation is expressed by negative values of correlation coefficient “rho” (ρ) with corresponding p-values.

## DISCUSSION

*In vivo* studies have shown changes in microbial communities due to rice bran or quercetin and describe the improvement of colonization resistance and reduction of colon cancer (11, 14, 53). However, these studies are marked by high variations of the microbiota composition likely due to host factors. Additionally, very little is known about the combined effect of rice bran and quercetin on gut microbiota. Thus, we focused on understanding the impact of rice bran extract and quercetin separately and in combination using a minibioractor array model(31). Also, to gain insights into taxonomical composition, we performed 16S rRNA sequencing over time and shotgun metagenome sequencing at the endpoint.

Using modified BHI medium, we were able to capture 30.93 % of the total amplicon sequence variants from the minibioreactors compared to the inoculum. This finding is slightly higher than previous results where the MBRA cultivated only 15-25% of the initial fecal community (28). The reason for the higher number of cultivated bacteria may be due to the modified BHI medium which has been shown to cultivate a wide range of gut bacteria (32). On the other hand, the highly dominant *Prevotellaceae* was lost in all four conditions in this study suggesting that fermenter adapted communities can lose much of their initial diversity (54). The alpha diversity indicated that the richness of the communities in bioreactors stabilized by day 14 (Supplementary Figure 1B). However, the taxonomical differences were evident as early as day four following inoculation. The changes in the relative abundances of taxa after day 14 were lower in control and quercetin supplemented medium whereas rice bran and combined rice bran with quercetin conditions had a more homogenized microbial composition after day 4 (Figure 2A). This indicates that the time required for the stabilization of communities may vary with the substrate used in minibioreactors in contrast to the previously determined timeframe of one week following fecal inoculation (28). Such changes in the time required for stabilization and diversity between the groups can be attributed ecologically to the stochastic rearrangement of the microbial species and their interactions at early stages because of selection pressure by substrate (55).

Rice bran is a nutrient-dense food with a unique profile and ratio of bioactive phytochemicals such as gamma oryzanol, tocotrienols, ferulic acid, vitamin B, beta-sitosterol and many others (12). It has been reported to be effective in preventing *Salmonella typhimurium* (9, 14, 56), rotavirus (57, 58) and norovirus (15) infections by priming intestinal immune cells. However, very few *in vivo* studies have highlighted its effect on gut microbiota and the reports have striking differences. In a clinical trial, Zambrana et al., (2019) showed significant enrichment of *Veillonella, Megasphaera* and *Dialister* species at the genus level from gut samples of children from either Nicaragua or Mali at 12 months’ time (24). Another study by Sheflin et al., (2015) showed a significant increase in the abundance of *Methanobrevibacter smithii, Paraprevotella clara, Ruminococcus flavefaciens, Dialister succinatiphilus, Bifidobacterium* sp., *Clostridium glycolicum, Barnesiella intestinihominis, Anaerostipes caccae* and *Ruminococcus bromii* OTUs after heat stabilized rice bran was fed to people (3 g/day) (25). The differences in the enriched taxa could be because of the unique inoculum used in this study. However, both above studies had lower resolution and report enrichment of different taxa which could be attributed to a different variety of rice bran being used and individualized host factors (59-61). The compounding hosts’ factors along with variation in age and geography (62), diet pattern (63), lifestyle (64-66), etc. play a crucial role in determining the gut microbiota composition, thus masking the actual effect of the substrate alone upon the diverse community. In contrast, this study supplies species-level resolution eliminating host interference and shows *Veillonella Prevotella, Dialister, Bacteroides* and *Alistipes* species are significantly enriched while *Oscillibacter, Eubacterium* and *Citrobacter* species are significantly reduced (Supplementary Figure 3B).

Similar to rice bran, previous studies analyzing microbial composition after quercetin supplementation yielded variable results among different hosts. *Enterobacteriaceae* and *Fusobacteriaceae* were reported to be positively related to quercetin supplementation whereas *Suttrellaceae* and *Oscillospiraceae* were found to be negatively correlated (26). However, Lei et al., (2019) reported an increase in abundances of *Bifidobacterium, Bacteroides, Lactobacillus* and *Clostridium*, with a reduction of *Fusobacterium* and *Enterococcus* in mice fed with quercetin (11). With higher resolution and removal of host factors, our results contrast with both studies and show enrichment of *Acidaminococcus intestini* and decreased abundances of *E. limosum, E. faecalis, A. caccae, D. longicatena* and *C. catus*. The domination of *Enterobacteriaceae* in medium supplemented with quercetin (Figure 2A, Supplementary Figure 3C) might have resulted in lower enrichment of the bacterial taxa as *Enterobacteriaceae* has been reported to affect quercetin metabolism by directly or indirectly inhibiting quercetin degrading bacteria (26).

When the combined effect of rice bran and quercetin were analyzed, we find that majority of the microbial shift is due to the supplementation of rice bran. (Figure 2, 3, Supplementary Figure 2). Most of the taxa enriched or decreased in the combination were very similar to those affected by rice bran supplementation alone. Compared to rice bran and quercetin supplementation separately, the combination was observed to significantly reduce members of the *Enterobacteriaceae* family (*Escherichia* and *Citrobacter* sp.) (Supplementary Table 1, Figure 6). *Enterobacteriaceae* consists of class of pathogens that are low in abundance but have potential to grow and dominate during dysbiotic conditions (67-69).The reduction of *Enterobacteriaceae* by combined rice bran and quercetin suggests a possibly beneficial effect on the host. The reduction of *Enterobacteriaceae* members was highly correlated with greater propionate levels in rice bran and quercetin combined medium (Figure 5G, Figure 6). The combined effect of rice bran and quercetin is reported for the first time in this study, where we show that the combination can alter the gut microbial composition and resulting metabolites to significantly reduce opportunistic pathogens in the gut

## CONCLUSION

Overall, this study provides evidence for species-level changes in the gut microbiota following quercetin and rice bran supplementation separately and in combination without host interference. Even though the impact of these dietary ingredients separately is described to be beneficial to human health, their combined effect was not known. Our results show that the combined effect of quercetin and rice bran reduces *Enterobacteriaceae* correlating to higher propionate levels. Thus, the combination of substrates such as rice bran and quercetin will be beneficial in excluding eneteric pathogens in the gut. However, it is crucial to culture, isolate and characterize the bacteria enriched in quercetin and rice bran supplemented medium to develop potential synbiotic formulations for gut health and immunity. Further, *in vivo* validation of such a combination would be necessary to understand its implication on the host.

## DATA AVAILABILITY

Raw sequence data from 16S rRNA amplicon sequencing and shotgun metagenome sequencing associated with this study had been deposited in NCBI Sequence Read Archive.

## ACKNOWLEDGMENTS

This work was supported in part by grants from the South Dakota Governor’s Office of Economic Development (SD-GOED) and the by the USDA National Institute of Food and Agriculture projects SD00H532-14 and SD00R646-18 awarded to J.S.

Computations supporting this project were performed on high-performance computing systems managed by the Research Computing Group, part of the Division of Technology and Security at South Dakota State University.

We declare no conflicts of interest.

---

**Supplementary Figure 1:**
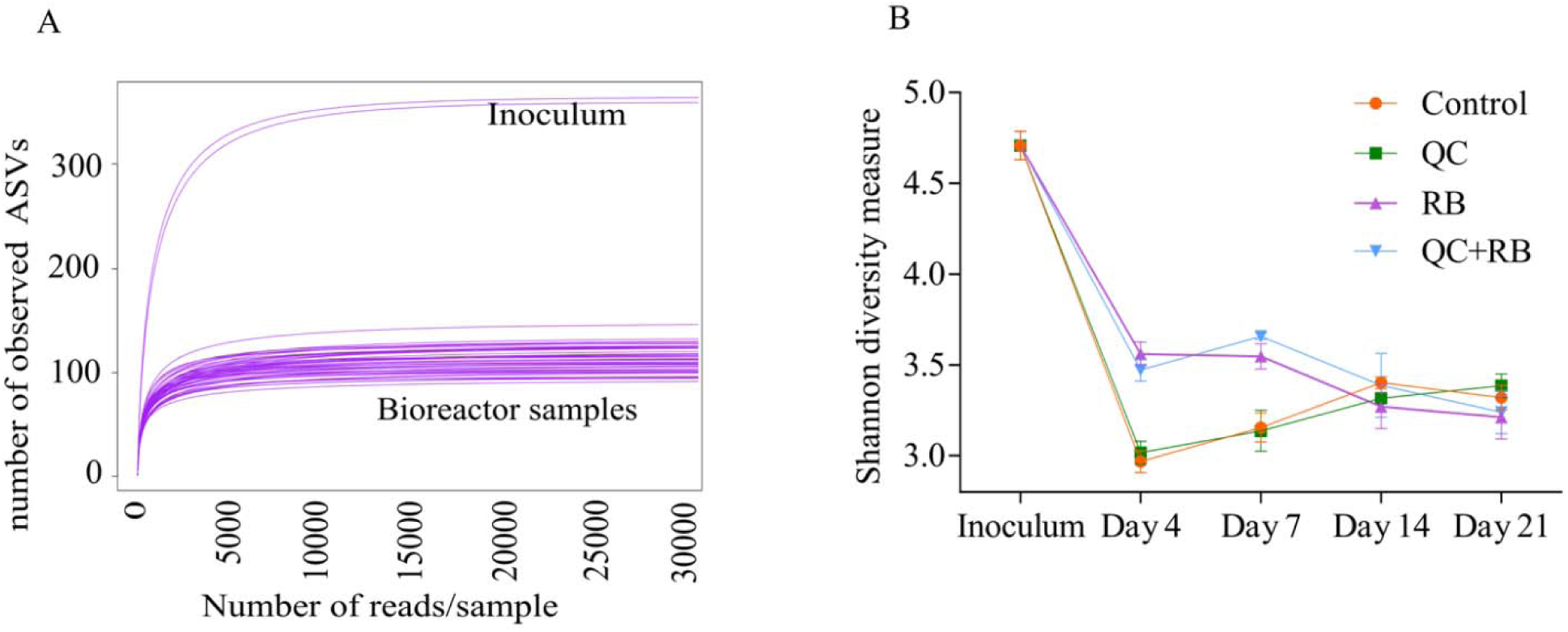
Exploratory analysis of predicted taxa and alpha diversity of microbiota in quercetin and rice bran: control, QC, RB and QC+RB medium and inoculum obtained using 16S rRNA community analysis. **A)** Rarefaction curves measuring the bacterial diversity in fecal communities. The curves are based on V3-V4 16S rRNA gene sequences obtained from a total of 48 samples (12 each from control, QC, RB, and QC+RB) from day 4, 7, 14 and 21 and duplicate inoculums **B)** Alpha diversity of four groups (control, QC, RB and QC+RB) at day 4, 7, 14 and 21.

**Supplementary Figure 2:**
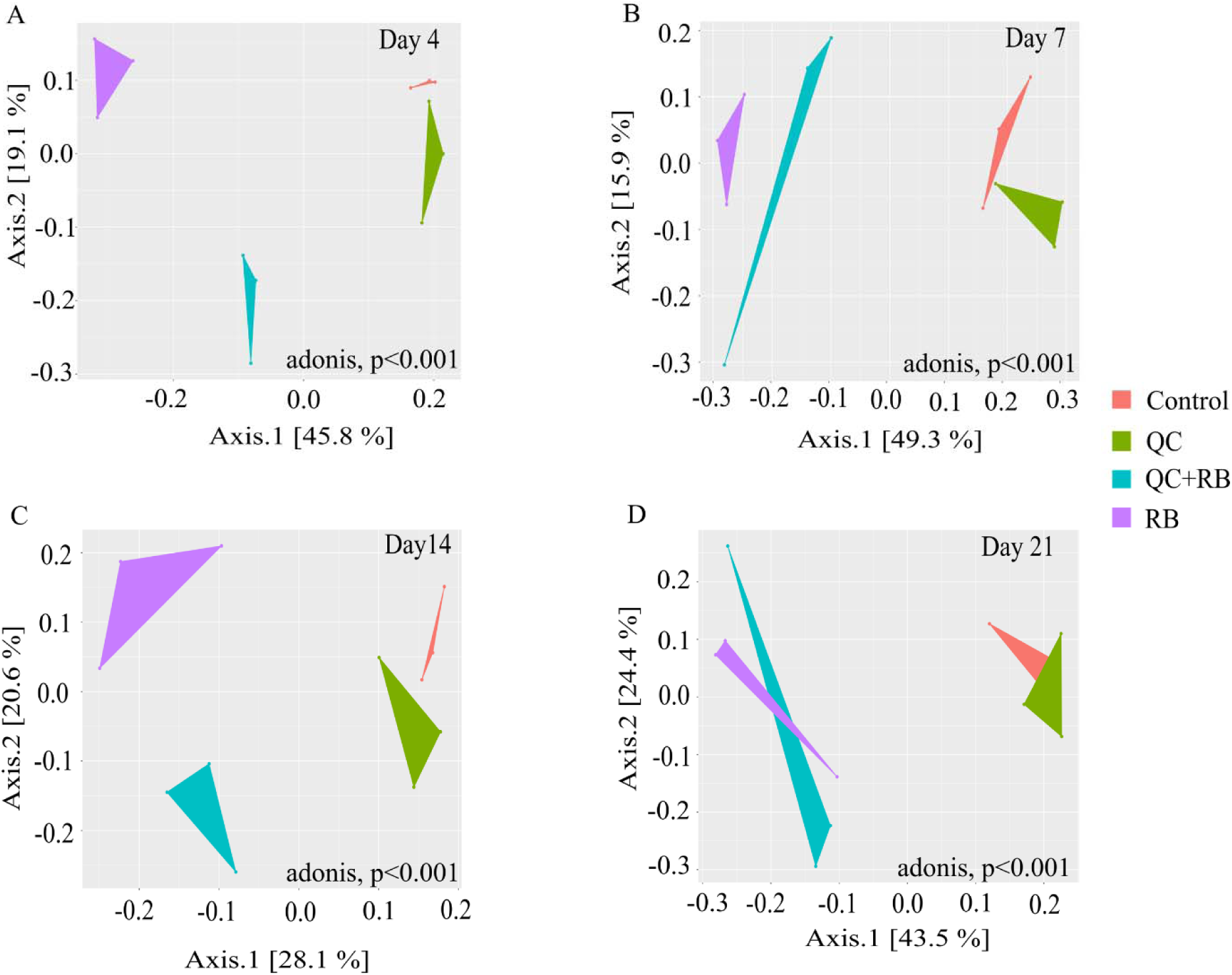
Beta diversity assessment of the microbiota communities formed at days 4, 7, 14 and 21 among four groups of media condition (Control, QC, RB and QC+RB) determined using 16S rRNA sequence analysis. The communities were ordinated using the “MDS” method followed by Bray-Curtis distance calculation to visualize the difference between the groups. Statistically, PERMANOVA using adonis was calculated on the beta diversity with p=0.004. The upper panel represents the beta diversity in early stages (day 4 and day 7) and the lower panel represents in later stages (day 14 and day 21).

**Supplementary Figure 3:**
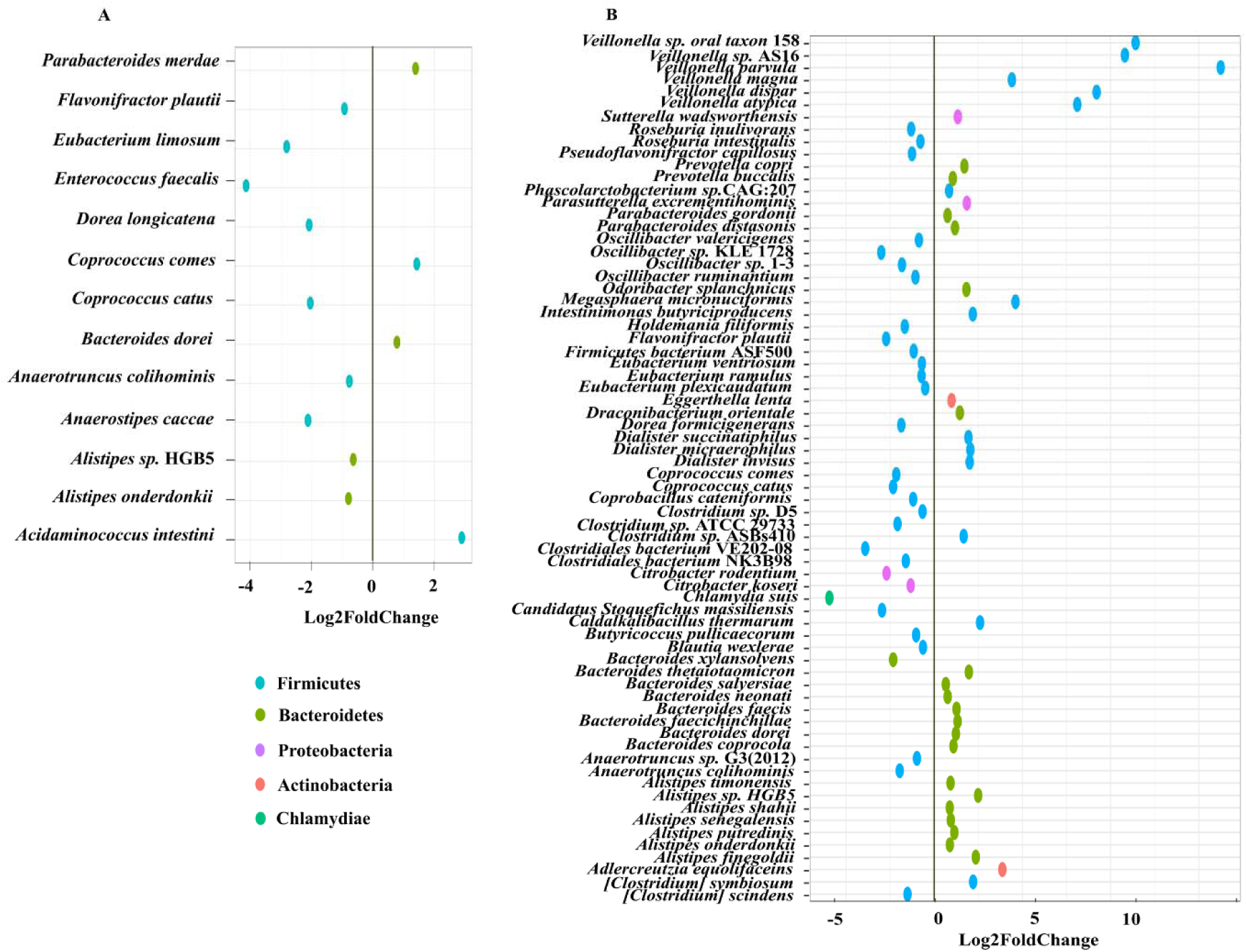
Impact of quercetin and rice bran (QC, RB) supplementation on bacterial species that are significantly altered in A) QC and B) RB when compared to the control medium. The endpoint (day21) shotgun metagenome sequence analysis was performed and the differential significance of the species was calculated by using DESEq2 (See Supplementary Table 1). The left column represents bacterial species-level taxa while the colors indicate their corresponding phyla (“Blue” =*Firmicutes*, “Brown” =*Bacteroidetes*, “Pink” =*Proteobacteria*, “Orange” =*Actinobacteria*, “Green” =*Chlamydiae*).

**Supplementary Figure 4:**
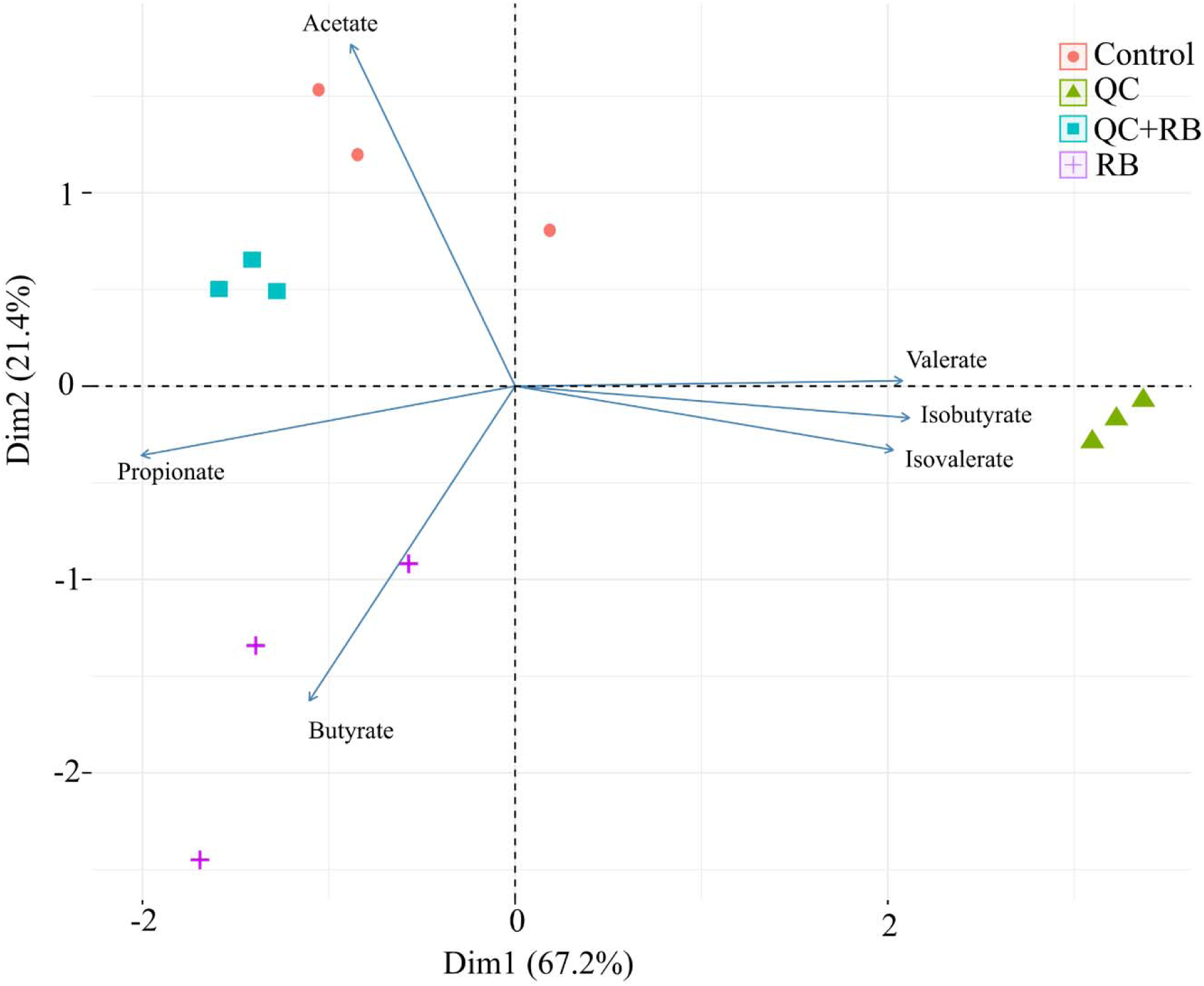
Biplot showing ordination of short-chain fatty acid production and bacterial communities formed at day 21 in control, QC, RB and QC+RB conditions. Abundances of the bacterial communities at day 21 determined by shotgun metagenomics were used with SCFAs profiles of day 21 to generate the plot.

**Supplementary Table 1:** List of all differentially abundant taxa in medium with quercetin (QC), medium with rice bran (RB) and medium with both quercetin and rice bran (QC+RB) compared to control. DESEq2 was used in shotgun metagenome sequences on day 21 for identification of differentially present taxa in the quercetin and rice bran supplemented media conditions when compared to control.

